# Translation of dipeptide repeat proteins in *C9ORF72*-ALS/FTD through unique and redundant AUG initiation codons

**DOI:** 10.1101/2022.08.06.503063

**Authors:** Yoshifumi Sonobe, Soojin Lee, Gopinath Krishnan, Yuanzheng Gu, Deborah Y. Kwon, Fen-Biao Gao, Raymond P. Roos, Paschalis Kratsios

## Abstract

A hexanucleotide repeat expansion in the first intron of *C9ORF72* is the most common monogenic cause of amyotrophic lateral sclerosis (ALS) and frontotemporal dementia (FTD). A hallmark of ALS/FTD pathology is the presence of dipeptide repeat (DPR) proteins, produced from both sense GGGGCC (poly-GA, poly-GP, poly-GR) and antisense CCCCGG (poly-PR, poly-PG, poly-PA) transcripts. Although initiation codons and regulatory factors have been identified for sense DPR translation, they remain mostly unknown for antisense DPRs. Here, we show that an AUG initiation codon is necessary for poly-PR synthesis, suggesting canonical AUG dependent translation. Remarkably, although an AUG located 194 base pairs (bp) upstream of the repeat is the main start codon for poly-PG synthesis, two other AUG codons (−212 bp, -113 bp) can also initiate translation, demonstrating a striking redundancy in start codon usage. eIF2D is required for CUG start codon-dependent poly-GA translation from the sense transcript in human motor neurons derived from induced pluripotent stem cells of *C9ORF72* ALS/FTD patients, but AUG-dependent poly-PG or poly-PR synthesis does not require eIF2D, indicating that distinct translation initiation factors control DPR synthesis from sense and antisense transcripts. Our findings provide key molecular insights into DPR synthesis from the *C9ORF72* locus, which may be broadly applicable to many other nucleotide-repeat expansion disorders.

## INTRODUCTION

The hexanucleotide GGGGCC repeat expansion in the first intron of *C9ORF72* is the most common monogenic cause of inherited amyotrophic lateral sclerosis (ALS) and frontotemporal dementia (FTD) ^1, 2^. This mutation is thought to cause ALS/FTD via three non-mutually exclusive mechanisms: (1) loss-of-function due to reduced C9ORF72 protein expression, toxicity from repeat-containing sense (GGGGCC) and antisense (CCCCGG) RNA^3, 4^, and (3) toxicity from dipeptide repeat (DPR) proteins produced from these transcripts^5^. DPRs produced from both sense (poly-GA, poly-GP, poly-GR) and antisense (poly-PR, poly-PG, poly-PA) transcripts are present in the central nervous system of ALS/FTD patients ^6, 7^. Strong evidence from experimental model systems suggests DPRs are toxic^8^, underscoring the importance of uncovering the molecular mechanisms responsible for DPR synthesis.

To design therapies that reduce DPR levels, it is valuable to identify initiation codons used in DPR translation. To date, the synthesis of sense DPRs has been a major focus in the ALS/FTD field, resulting in the identification of translation initiation codons for poly-GA and poly-GR ^9, 10, 11, 12^. As previously shown, *non-canonical* codons (viz., CUG for poly-GA, AGG for poly-GR) initiate DPR synthesis from the sense strand, suggesting an unconventional form of translation, i.e., repeat-associated non-AUG (RAN) translation^6^. However, deletion analysis of *cis*-regulatory elements upstream of the GGGGCC repeats and ribosome profiling revealed that translation of the poly-GA and poly-GR frames is independent of the presence of G_4_C_2_ repeats^13, 14, 15^. Moreover, a recent study reported that a canonical AUG initiation codon (194 nucleotides upstream of the repeat) is used for poly-PG synthesis from the antisense CCCCGG transcript, suggesting conventional translation is involved in the synthesis of at least one DPR. Despite the latter finding, the initiation codon for other DPRs (e.g., poly-PR) from the antisense transcript remains unknown. Hence, it is unclear which form of translation (RAN vs. conventional) is utilized for DPR synthesis from the antisense transcript. Studying the mechanisms responsible for DPR synthesis from the antisense transcript is important because a recent ALS clinical trial that specifically targeted the production of sense DPRs failed. In the latter case, no improvements in clinical outcomes occurred despite decreased levels of sense DPRs ^16, 17^.

An additional challenge in ALS/FTD is the identification of regulatory factors necessary for DPR synthesis. Research efforts have uncovered a number of proteins that act at different steps of DPR synthesis: RNA helicase eIF4A^9^, cap-binding initiating factor eIF4E^18^, small ribosomal protein subunit 25 (RPS25)^19^, eukaryotic translation initiation factors eIF2A^12^, eIF3F^20^, eIF2D^21^, and eIF2D co-factors DENR and MCTS-1^22^. Except for RPS25, all remaining factors have only been assessed for their effects on DPRs produced from the sense GGGGCC transcript. Hence, it remains unknown whether any of these factors is used for DPR synthesis from the antisense transcript. Furthermore, the role of these factors on DPR synthesis in induced pluripotent stem cell (iPSC)-derived neurons from *C9ORF72* ALS/FTD patients remains largely untested.

Here, we employ cell-based models of *C9ORF72* to identify translation initiation codons for DPRs produced from the antisense transcript. We find that a canonical AUG initiation codon located 273 base pairs (−273 bp) upstream of the CCCCGG repeats is necessary for poly-PR synthesis. Furthermore, we provide evidence for redundancy in usage of canonical initiation codons for poly-PG synthesis, as follows. Although an AUG at -194 bp is the main start codon for poly-PG, two other AUG codons (at -212 bp and at -113 bp) can also function as translation initiation sites. These findings suggest that DPR synthesis from the antisense transcript occurs via AUG dependent translation, contrasting with the DPR synthesis from the sense transcript, which depends on near-cognate start codons (CUG for poly-GA, AGG for poly-GR). Furthermore, we critically extend previous observations made in *C. elegans* and cell-based models^21^ by demonstrating that translation initiation factor eIF2D is necessary for CUG-dependent poly-GA synthesis from the sense transcript in iPSC-derived motor neurons from *C9ORF72*-ALS/FTD patients. However, eIF2D is not involved in AUG-dependent antisense DPR (poly-PG, poly-PR) synthesis, suggesting that translation initiation sites and factors for DPR synthesis from sense GGGGCC and antisense CCCCGG transcripts are distinct.

## RESULTS

### A canonical AUG initiation codon located 273 bp upstream of CCCCGG repeats is required for poly-PR synthesis

To study DPR synthesis from the antisense transcript, we engineered three constructs with 35 CCCCGG repeats preceded by 1000bp-long intronic sequence from human *C9ORF72* (**Fig. 1 A**) ^21^, and then followed by nanoluciferase (NanoLuc) in frame of poly-PR, poly-PG, or poly-PA (see Materials and Methods). Upon transfection into HEK293 or NSC34 cells, robust expression of poly-PR and poly-PG, but not poly-PA, was present in luciferase assays (**Fig. 1B-C**) and Western blots (**Fig. 1D-E**). These poly-PR::NanoLuc and poly-PG::NanoLuc constructs offer an opportunity to identify the initiation codons for poly-PR and poly-PG synthesis.

**Figure 1.**
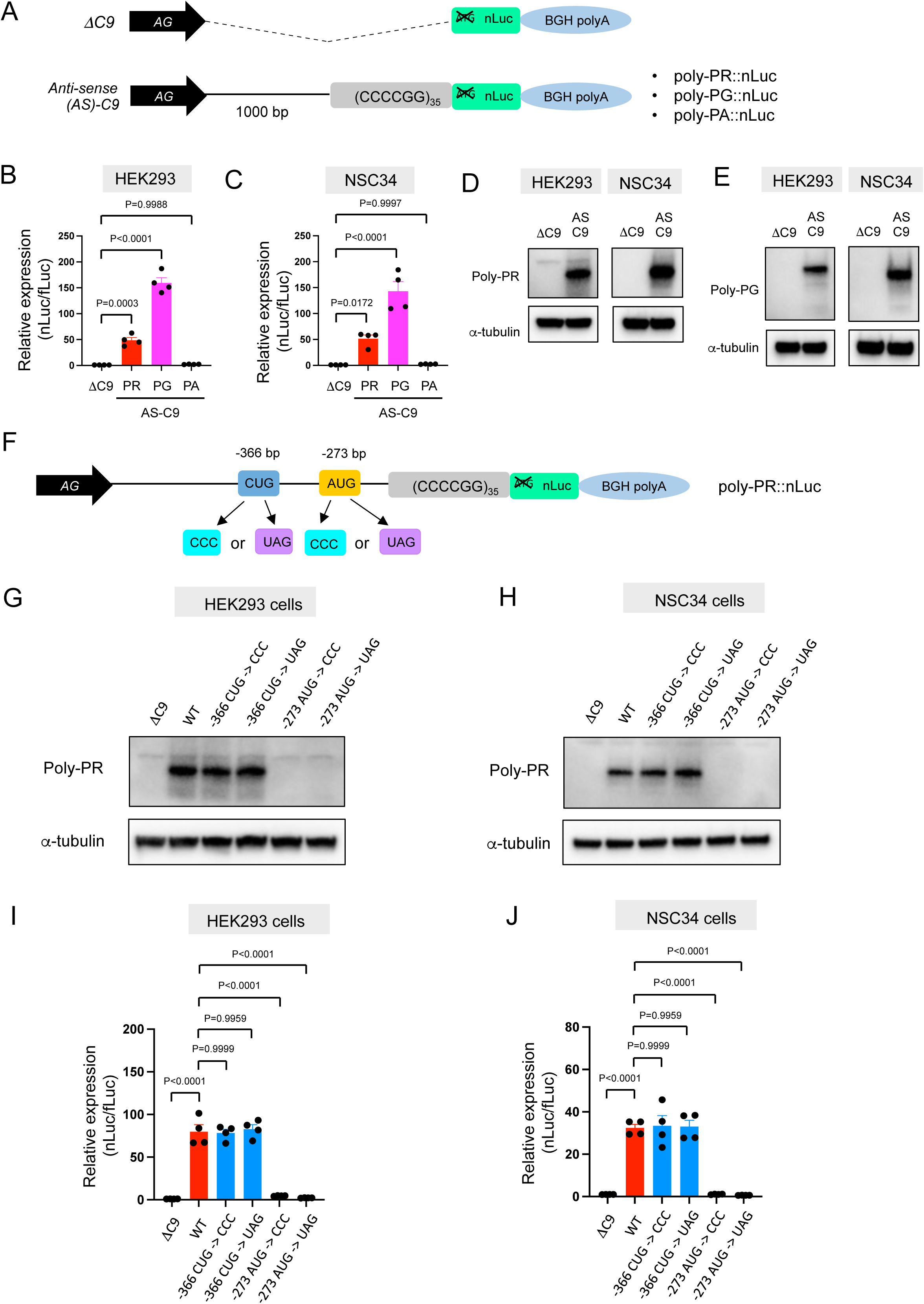
Poly-PR and poly-PG are translated from antisense CCCCGG repeats. (A) Schematic diagram of the constructs with 35 CCCCGG repeats preceded by 1000bp-long intronic sequence from human *C9ORF72*, and then followed by nanoluciferase (nLuc). (B-C) (B) HEK293 and (C) NSC34 cells were cotransfected with fLuc along with either ΔC9 or AS-C9 plasmids. The levels of luciferase activity were assessed by dual luciferase assays (mean ± s.e.m.). One-way ANOVA with Tukey’s multiple comparison test was performed. (D-E) HEK293 and NSC34 cells were transfected with either ΔC9 or AS-C9 plasmids. Cell lysates were processed for Western blotting, and immunostained with antibodies to (D) poly-PR, (E) poly-PG, and α-tubulin. The experiments were repeated 4 times. (F) Schematic diagram showing the mutants of putative start codons for poly-PR. (G) HEK293 and NSC34 cells were transfected with the indicated plasmids. Cell lysates were processed for Western blotting, and immunostained with antibodies to poly-PR and α -tubulin. (H-I) HEK293 and NSC34 cells were cotransfected with the plasmids along with fLuc. The level of luciferase activity was assessed by dual luciferase assays (mean ± s.e.m.). One-way ANOVA with Tukey’s multiple comparison test was performed. The experiments were repeated 4 times.

We initially focused on poly-PR, one of the most toxic DPRs based on *in vitro* ^23, 24, 25^ and *in vivo* studies in worms ^26^, flies ^23, 27, 28^ and mice ^28, 29, 30^. Using our recently developed machine-learning algorithm for initiation codon prediction ^31^, we identified a CUG at -366bp (Kozak sequence: guaCUGa) and an AUG at -273bp (Kozak sequence: cggAUGc) as putative initiation codons for poly-PR (**Fig. 1F**). We then mutated these codons either to CCC or the termination codon UAG (**Fig. 1F**). Western blotting and luciferase assay showed that mutation of the CUG at -366bp to CCC or UAG did not affect poly-PR expression (**Fig. 1G-J**). However, mutation of the AUG at -273bp to CCC or UAG completely abolished poly-PR expression both in HEK293 and NSC34 cells (**Fig. 1G-J**). These results strongly suggest that AUG at -273bp is the start codon for poly-PR. Of note, a previous study also detected poly-PR synthesis when only 100 bp of intronic sequence downstream of the GGGGCC repeats was cloned in an adeno-associated viral vector^32^. Although the intronic sequence was only 100 bp-long, it was located next to a 589 bp regulatory element of the woodchuck hepatitis virus (WPRE) that contains several putative start codons for poly-PR synthesis.

### Evidence for redundancy of AUG initiation codon usage in poly-PG translation

We next investigated poly-PG, which is less toxic than poly-PR ^23, 27, 33, 34^, and has been proposed as a biomarker for *C9ORF72*-ALS/FTD ^35, 36^. Using the same machine-learning algorithm ^31^, we identified four putative initiation codons (AUG at -212bp, AUG at -194bp, CUG at -182bp, AUG at -113bp) (**Fig. 2A**), all with relatively good Kozak sequences (gaaAUGa at -212bp, aaaAUGc at -194bp, gctCUGa at -182bp, aggAUGc at -113bp). Of note, a prior publication previously identified the AUG at -194bp as an initiation codon ^11^. Mutation of all four of these codons to CCC completely blocked poly-PG expression (**Fig. 2B-D**), suggesting one or more of these codons is required. Next, we simultaneously mutated three codons to CCC, but left intact the AUG at -212bp. As a result of this change, we observed poly-PG expression, suggesting poly-PG translation can start at the AUG at - 212bp. Intriguingly, when we followed a similar approach to mutate 3 codons to CCC but leave intact the AUG at -194bp or at -113bp, we also observed poly-PG production, but this time at an expected lower molecular weight (**Fig. 2B-D**). Of note, when we mutated to CCC all three AUG codons (−212bp, -194bp, -113bp) but left intact the CUG at -182bp, we observed no poly-PG expression (**Fig. 2B-D**). These results suggest that any of these three AUGs, but not the CUG at -182bp, can function as a start codon for poly-PG, indicating redundancy in the translation initiation codon for poly-PG.

**Figure 2.**
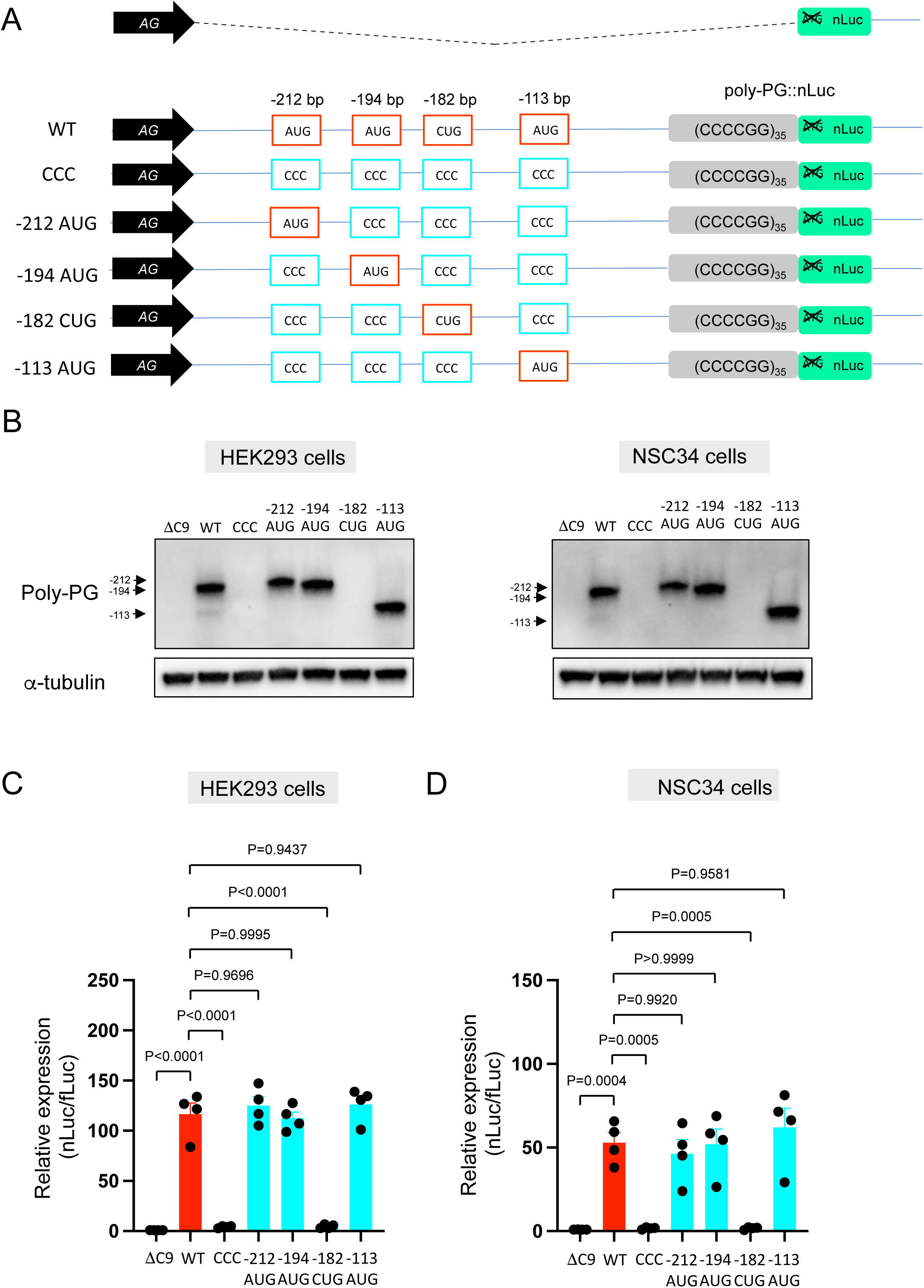
An AUG at -194bp position is the start codon for poly-PG translation. Schematic diagram showing mutants with changes in the putative start codons for poly-PG. (B) HEK293 and NSC34 cells were transfected with indicated plasmids. Cell lysates were processed for Western blotting, and immunostained with antibodies to poly-PG and α-tubulin. (C-D) (C) HEK293 and (D) NSC34 cells were cotransfected with fLuc plasmid along with other indicated plasmids. The level of luciferase activity was assessed by dual luciferase assay. One-way ANOVA with Tukey’s multiple comparison test was performed. The experiments were repeated 4 times. mean ± s.e.m.

We observed a strong (higher molecular weight) band and a fainter (lower molecular weight) band for poly-PG when the intact version of the poly-PG::NanoLuc plasmid was translated (**Fig. 2B**). The strong band is likely to result from translation initiation at the AUG at -194bp, whereas the faint band is likely initiated at the AUG at -113bp (**Fig. 2B**). Hence, the AUG at -194bp appears to be the main initiation codon for poly-PG synthesis from the antisense transcript of 35 C4G2 repeats (**Fig. 2B**), which is consistent with mass-spectrometry results from a previous report ^11^. Of note, selective mutation of the AUG at -194 to CCC did not abolish poly-PG expression (**Fig. 3A-D**). Instead, it led to the production of two poly-PG products: a high molecular weight product (strong band) resulting from use of the AUG at -212bp as well as a lower molecular weight product (faint band) resulting from AUG at -113bp (**Fig. 3B**). Altogether, these results suggest that the AUG at -194bp is mainly used for poly-PG expression from antisense C4G2 repeats. However, when this AUG is mutated, two other AUG codons (at -212bp and -113bp) can also function as translation initiation sites, again revealing redundancy in the start codon usage for poly-PG synthesis.

**Figure 3.**
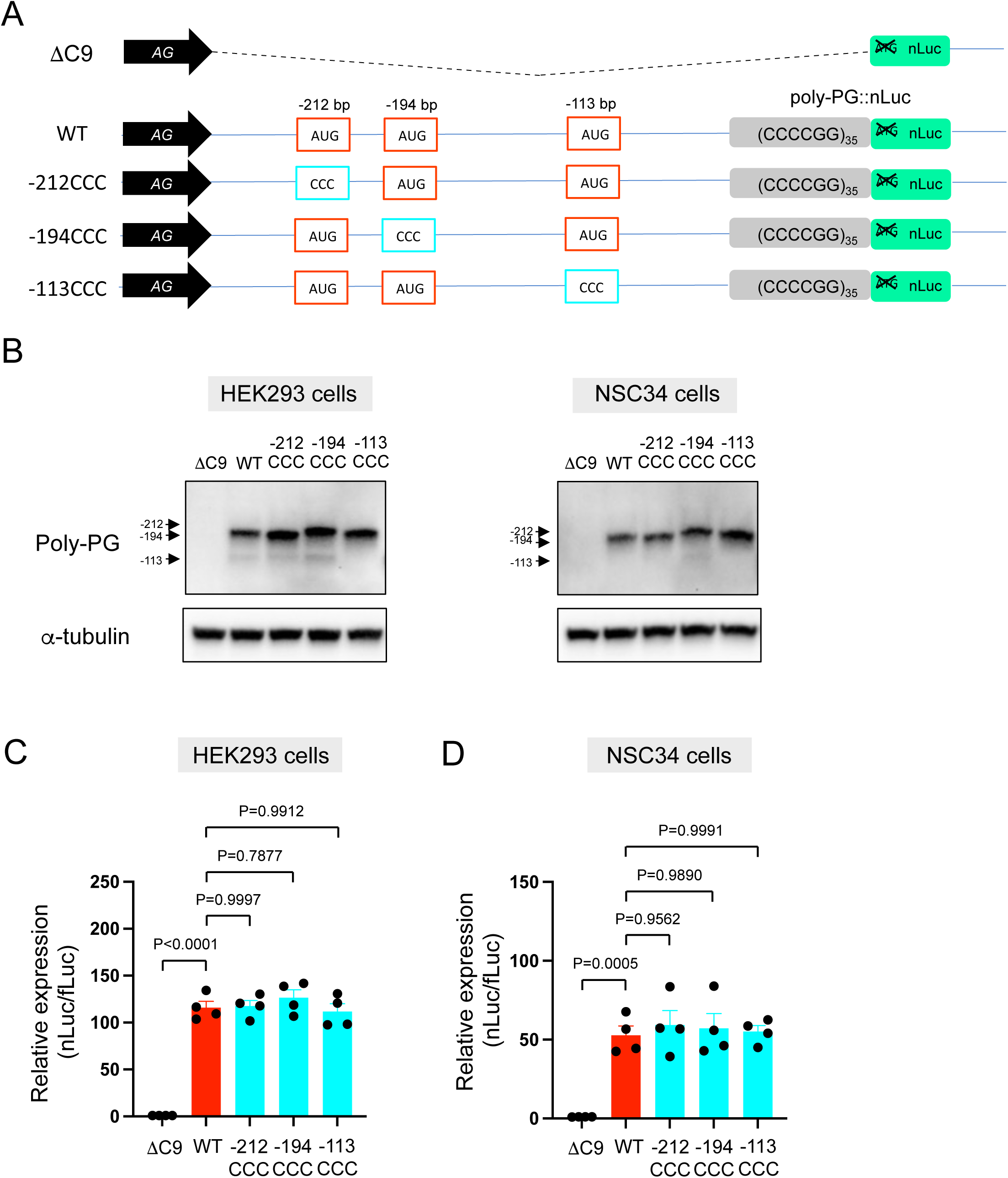
Mutation of AUG codons to CCC fails to suppress poly-PG translation. (A) Schematic diagram of the constructs. (B) HEK293 and NSC34 cells were transfected with indicated plasmids. Cell lysates were processed for Western blotting, and immunostained with antibodies to poly-PG and α-tubulin. (C, D) (C) HEK293 and (D) NSC34 cells were cotransfected with fLuc plasmid along with indicated plasmids. The level of luciferase activity was assessed by dual luciferase assays. One-way ANOVA with Tukey’s multiple comparison test was performed. The experiments were repeated 4 times. mean ± s.e.m.

We further corroborated redundant initiation for poly-PG translation by separately mutating each of the AUG codons to a termination UAG codon (**Fig. 4A-D**). Mutation of the AUG at -212bp to UAG failed to affect poly-PG expression, most likely because the AUG at -194bp became the start codon as shown by Western blots (**Fig. 4B-D**). Similarly, mutation of the AUG at -194bp to UAG did not affect poly-PG expression because the AUG at -113bp became the start codon (**Fig. 4B-D**). However, mutation of AUG at -113bp to UAG completely blocked poly-PG expression (**Fig. 4B-D**). Altogether, these findings strongly suggest that the AUG at -194bp is primarily used for poly-GP translation, but the other two AUG codons at -212bp and -113bp can also function as translation initiation sites under certain experimental conditions.

**Figure 4.**
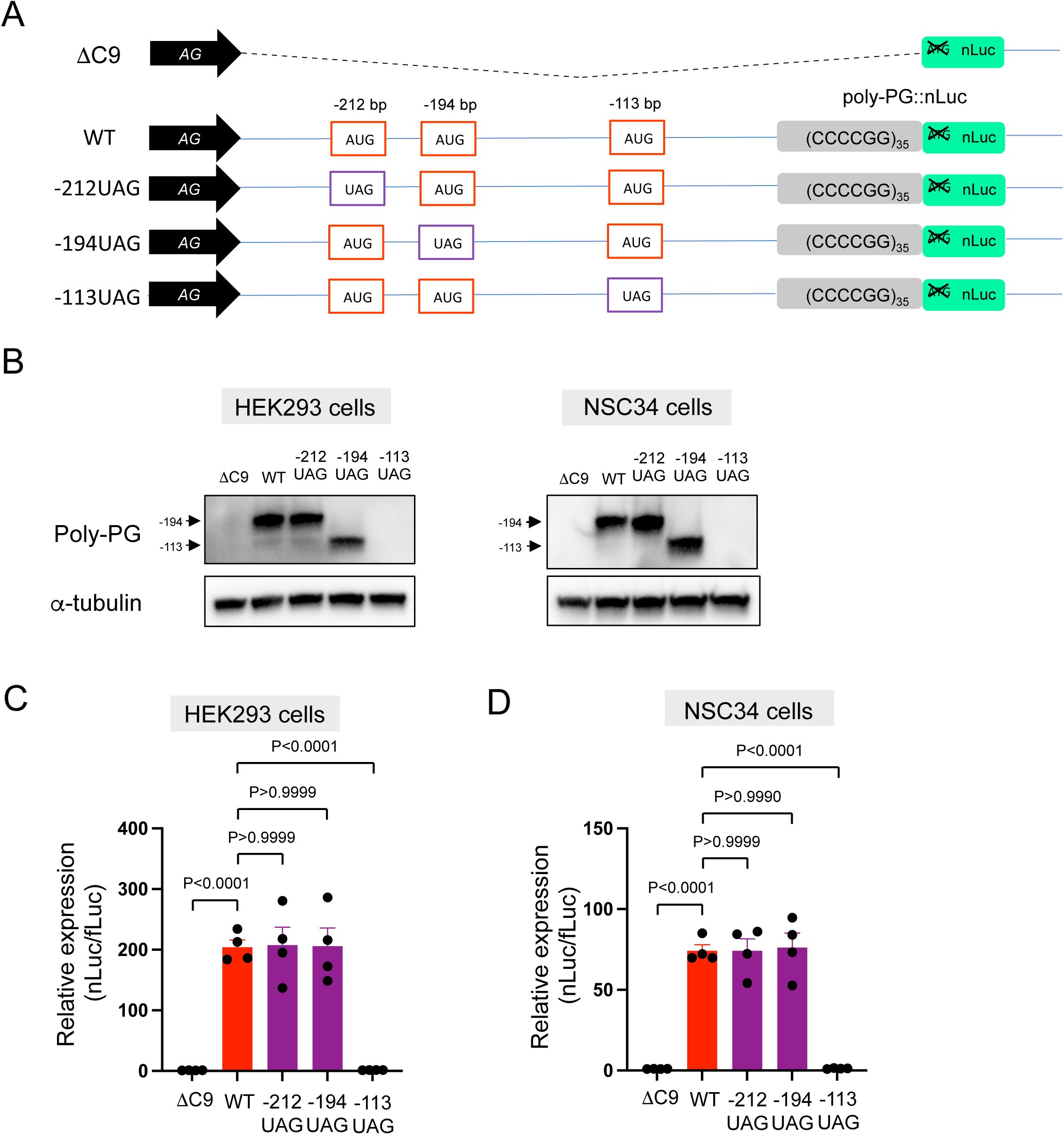
Redundancy of start codon usage in poly-PG translation. (A) Schematic diagram of the constructs. (B) HEK293 and NSC34 cells were transfected with indicated plasmids. Cell lysates were processed for Western blotting, and immunostained with antibodies to poly-PG and α-tubulin. (C, D) (C) HEK293 and (D) NSC34 cells were cotransfected with fLuc plasmid along with indicated plasmids. The level of luciferase activity was assessed by dual luciferase assays. One-way ANOVA with Tukey’s multiple comparison test was performed. The experiments were repeated 4 times. mean ± s.e.m.

### Knockdown of eIF2D does not affect poly-PG synthesis but reduces poly-GA in iPSC-derived motor neurons

Following the identification of AUG codons for translation initiation of poly-PG, we next sought to identify translation initiation factors necessary for this DPR synthesis. We focused on eIF2D, since we had previously found it to be necessary for poly-GA synthesis from the sense transcript in *C. elegans* and cell-based models (HEK293 and NSC34 cell lines) ^21^. To test whether eIF2D has a role in poly-PG translation, we used a published iPSC line from a *C9ORF72* carrier, as well as its isogenic control line which had CRISPR/Cas9-mediated deletion of expanded GGGGCC repeats^37^. The iPSC lines were differentiated into motor neurons as previously described^38^. Repeated transfection of a small interfering RNA (siRNA) against eIF2D, but not a control scrambled siRNA, resulted in robust downregulation of *eIF2D* mRNA as assessed by RT-PCR (**Fig. 5A**). The mRNA levels of eIF2A, a related initiation factor, remained unaltered, suggesting specificity in the siRNA effect. Despite this knockdown, an immunoassay failed to show any differences in the steady-state levels of poly-PG (**Fig. 5B**), suggesting eIF2D is not necessary for poly-PG translation from the antisense transcript. Although this assay does not distinguish between poly-PG produced from the antisense transcript and poly-GP from the sense transcript, PG/GP inclusions in brain tissue of *C9ORF72* ALS/FTD patients contain ∼80% of poly-PG from the antisense transcript and ∼20% of poly-GP from the sense transcript^6^. Hence, our data suggest that eIF2D does not affect poly-PG synthesis from the antisense CCCCGG transcript.

**Figure 5.**
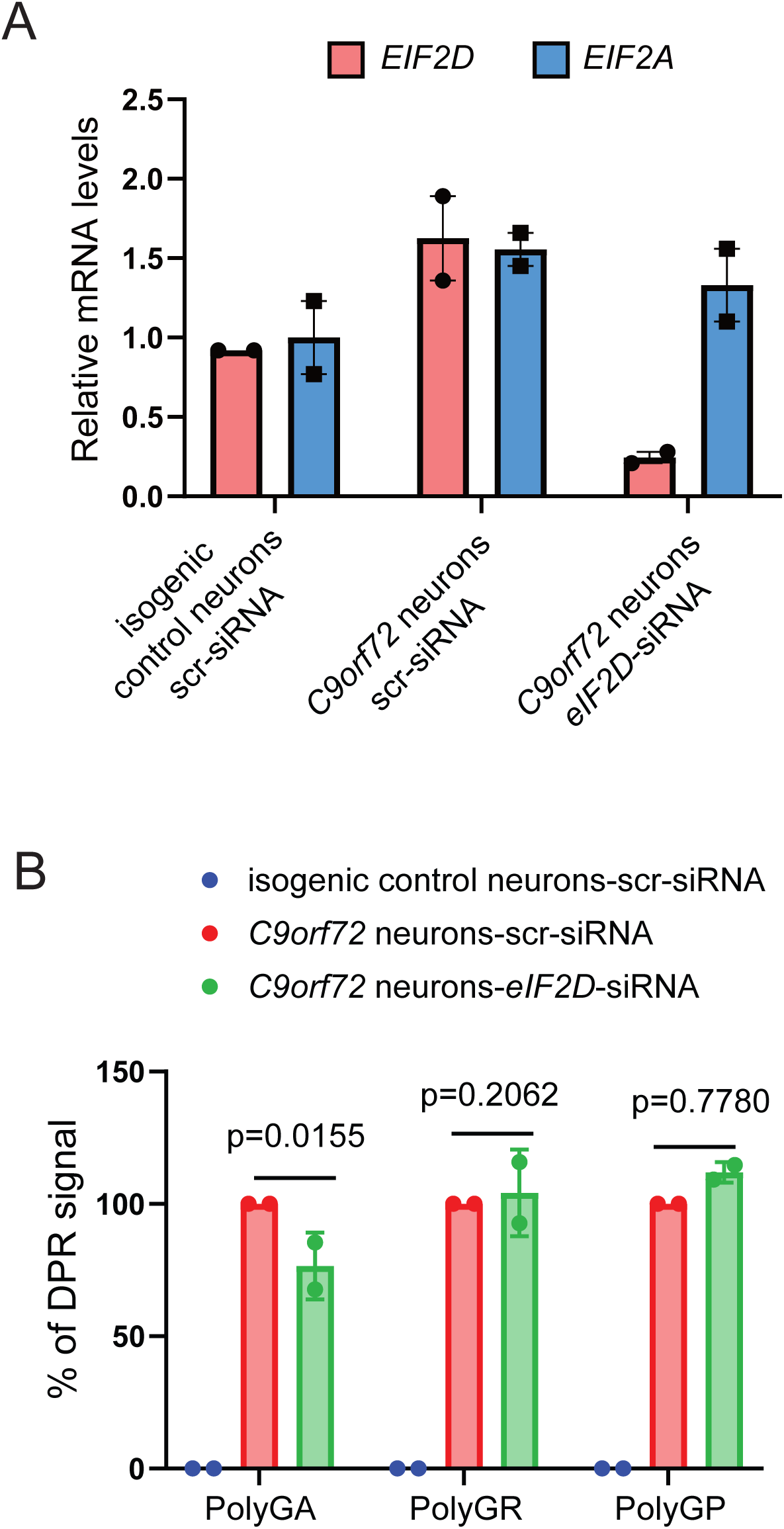
Knockdown of eIF2D reduces poly-GA steady-state levels in human iPSC-derived neurons. (A) The *eIF2D*, *eIF2A*, and *actin* mRNA levels were assessed by real-time quantitative PCR on either isogenic control or *C9ORF72* human motor neurons upon siRNA transfection (scramble or EIF2D siRNA). The *eIF2D* and *eIF2A* mRNA levels were normalized to actin. The experiments were repeated two times. P<0.05 by Two-tailed unpaired *t*-test. (B) Poly-GA, poly-GR and poly-GP levels in motor neurons differentiated independently (twice) from isogenic control and *C9ORF72* iPSC lines. DPR levels were measured using an MSD immunoassay. Data presented as mean ± S.D. *P* values were calculated using 2-way ANOVA with Dunnett’s multiple comparison test using Prizm (9.1) software.

Despite the above findings, eIF2D knockdown significantly affects poly-GA synthesis from the sense GGGGCC transcript in iPSC-derived neurons, critically extending previous observations made in *C. elegans* and cell-based models ^21^ (**Fig. 5B**). Consistent with the latter study, eIF2D knockdown had no effect on poly-GR synthesis from the sense transcript (**Fig. 5B**). Altogether, these findings suggest that eIF2D is required for CUG start codon dependent poly-GA synthesis from the sense transcript in human iPSC-derived neurons, but is dispensable for poly-GR and poly-PG synthesis from sense and antisense transcripts, respectively.

### eIF2D does not control poly-PR and poly-PG synthesis from the antisense transcript

Since immunoassays to measure poly-PR steady-state levels in human iPSC-derived neurons are not yet established, we transfected the poly-PR::NanoLuc reporter construct into HEK293 in order to evaluate the effect of eIF2D in poly-PR synthesis. To this end, we generated an *EIF2D* knockout HEK293 line using CRISPR/Cas9 gene editing (see Materials and Methods), and then performed a luciferase assay to measure poly-PR::NanoLuc expression (**Fig. 6A-B**). We found that knockout of *EIF2D* did not affect the expression levels of poly-PR (**Fig. 6C**). Importantly, we obtained similar results upon knockdown of *EIF2D* with a short hairpin RNA (shRNA) (**Fig. 6D**), again suggesting that eIF2D is not required for poly-PR synthesis from antisense CCCGG transcripts. Lastly, knock-out (CRISPR/Cas9) or knock-down (shRNA) of *eIF2D* in HEK293 cells had no effect on poly-PG::NanoLuc reporter expression (**Fig. 6C-D**), corroborating our findings in human iPSC-derived neurons (**Fig. 5**).

**Figure 6.**
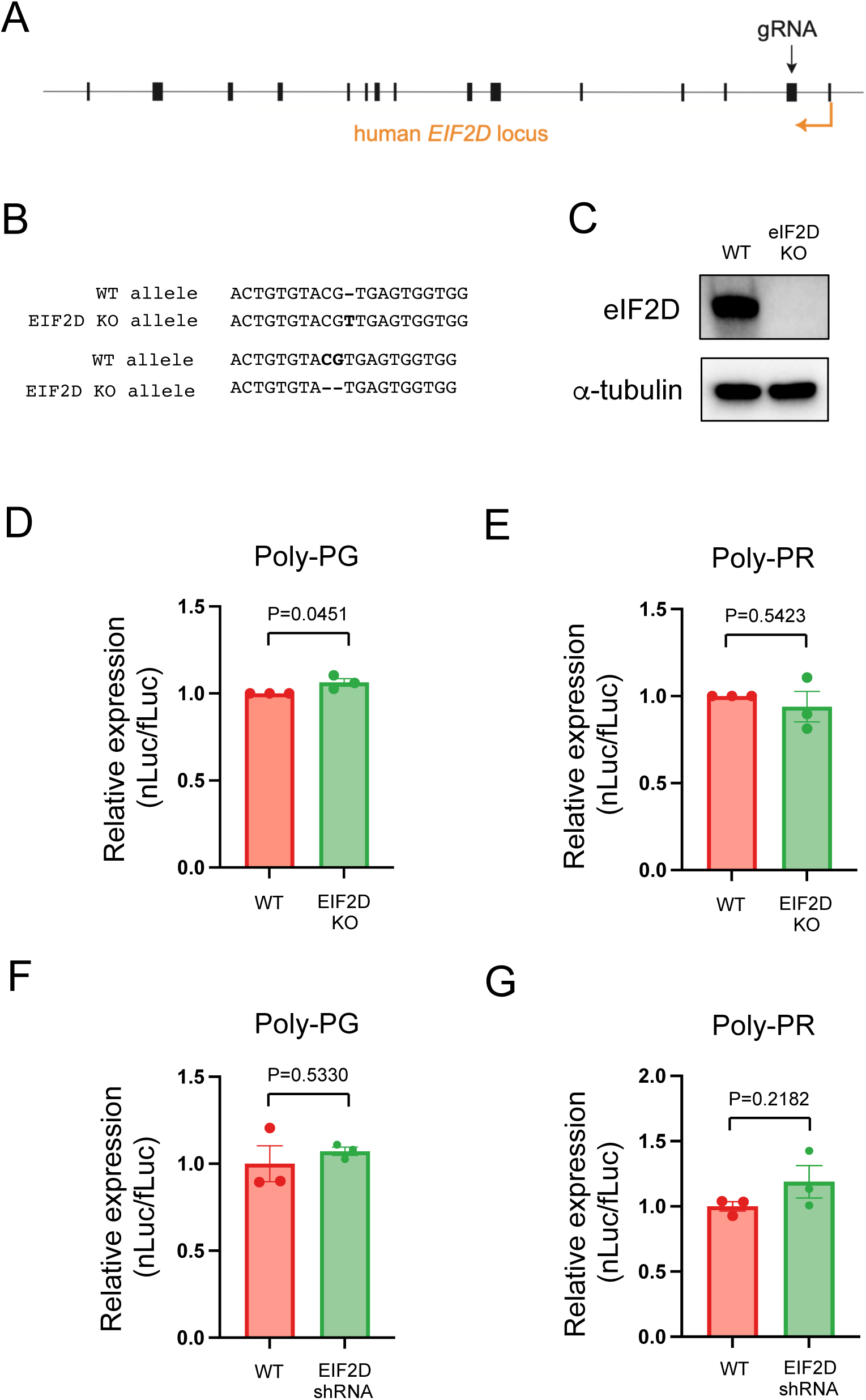
Downregulation of *EIF2D* does not reduce expression levels of poly-PG and poly-PR. (A) A gRNA targeted the second exon of human EIF2D (see Materials and Methods). (B) After CRISPR/Cas9-mediated gene editing, the *EIF2D* knockout (EIF2DKO) HEK293 cells carried different mutations on each allele. (C) Cell lysates from WT and EIF2DKO HEK293 cells were processed for Western blotting, and immunostained with antibodies to eIF2D and α-tubulin. (D-E) WT and EIF2DKO HEK293 cells were cotransfected with fLuc plasmid along with AS-C9 plasmids. The level of luciferase activity was assessed by dual luciferase assays. (F-G) WT HEK293 cells were transfected with fLuc and AS-C9 plasmids along with anti-EIF2D shRNA. The level of luciferase activity was assessed by dual luciferase assays. Unpaired t test was performed. N = 3. mean ± s.e.m.

## DISCUSSION

Here, we show that canonical AUG codons on the antisense CCCCGG transcript serve as translation initiation codons for two DPRs, viz., poly-PR and poly-PG. This finding may inform the design of future therapy for ALS/FTD, especially since poly-PR is a highly toxic DPR and poly-PG synthesis is primarily translated from the antisense transcript ^6^. Our finding of canonical AUG codons serving as translation initiation codons for antisense DPRs (poly-PR, poly-PG) differs from the proposed mode of translation of sense DPRs (poly-GA, poly-GR). In the latter case, it is thought that repeat-associated non-AUG (RAN) translation of poly-GA and poly-GR occurs via non-canonical CUG and AGG codons, respectively ^9, 10, 11, 12, 14, 21^. However, this model of RAN translation for poly-GA and poly-GR has been recently challenged, as translation of these DPRs does not depend on the presence of GGGGCC repeats^13, 14, 15, 21^. Nevertheless, our findings merged with those of previous studies suggest that DPR synthesis involves at least two different modes of translation: near-cognate start codon (e.g., CUG, AGG) dependent translation for poly-GA and poly-GR from sense GGGGCC transcripts, as well as conventional AUG dependent translation for poly-PR and poly-PG synthesis from antisense CCCCGG transcripts.

A notable finding of the present study is the presence of redundancy in start codon usage for poly-PG synthesis under specific experimental conditions. Our findings suggest that the AUG at - 194bp is primarily used for poly-GP translation from antisense CCCCGG transcripts, consistent with a previous investigation^11^. However, when this AUG is mutated, two other canonical AUG codons (at - 212bp and -113bp can also function as translation initiation sites under certain experimental conditions. Although it remains unknown whether such redundancy of translation initiation occurs in the central nervous system of *C9ORF72* ALS/FTD patients, these findings nevertheless suggest that targeting only one translation initiation site may be insufficient to prevent poly-PG synthesis. We note that redundancy in start codon usage may also apply to poly-PR synthesis from the antisense transcript: although we identified an AUG at -273 bp as necessary for poly-PR synthesis, a previous study detected poly-PR when only 100bp downstream of the GGGGCC repeats were included in an adeno-associated viral vector^32^.

Emerging evidence suggests distinct mechanisms affect translation initiation of DPRs from sense and antisense transcripts in *C9ORF72* ALS/FTD. For example, the RNA helicase DDX3X directly binds to sense (GGGGCC), but not antisense (CCCCGG) transcripts, thereby selectively repressing the production of sense DPRs (poly-GA, poly-GP, poly-GR)^39^. Further, the accessory proteins eIF4B and eIF4H interact directly with sense GGGGCC transcripts and are required for poly-GR synthesis in a *Drosophila* model of *C9ORF72* ALS/FTD^40^. Here, we provide evidence that the translation initiation factor eIF2D is not involved in DPR (viz., poly-PG, poly-PR) synthesis from antisense (CCCCGG) transcripts, but is selectively required for poly-GA production from sense (GGGGCC) transcripts in human iPSC-derived motor neurons. The latter findings are important because they indicate that distinct initiation sites and factors are involved in DPR translation from sense and antisense transcripts, perhaps a reflection of the different modes of translation (RAN- and AUG-dependent translation) of DPRs. Consistent with the idea of distinct factors being involved, translation initiation is the most heavily regulated step in protein synthesis because it is the rate-limiting step of this process^41^. In contrast to the different mechanisms responsible for DPR translation, the transcriptional control of sense and antisense transcripts appears coordinated. For example, a single protein – the transcription elongation factor Spt4 – controls production of both sense and antisense transcripts^42^.

In addition to *C9ORF72*-ALS/FTD, nucleotide repeat expansions are present in various genes, causing more than 30 neurogenetic diseases^43, 44^. In many of these disorders, products translated from the expanded repeat sequences have been detected in the nervous system of affected individuals. The findings of the present study may also apply to this large group of genetic disorders in the following ways. First, translation of peptides from the same nucleotide repeat expansion may require different modes of translation (RAN- and AUG-dependent translation), as previously proposed^45^. Second, the surprising redundancy in canonical AUG codon usage for poly-PG may also apply to proteins translated from nucleotide repeat expansions in other genes, as the number of nucleotide repeats is often variable in different neural cells of the same patient. Lastly, our results support the idea that distinct translation initiation factors are involved in the synthesis of individual DPRs produced from the same nucleotide repeat expansion. This finding suggests that the design of therapies for diseases caused by expanded nucleotide repeats may be especially challenging.

## Acknowledgements

This work was supported by a grant from the Lohengrin Foundation (R.P.R, P.K), a basic science pilot grant from the Association for Frontotemporal Degeneration (AFTD) (R.P.R, P.K), and two NIH grants (R37NS057553 and R01NS101986) to F.B.G. The antibodies used to measure GA levels were discovered by Neurimmmune AG (Zurich, Switzerland)”.

## Author Contributions

Study design: Y.S., R.P.R., P.K. Literature search: Y.S., R.P.R., P.K. Experimental studies: Y.S., S.L., G.K., Y.G., D.Y.K. Data analysis/interpretation: Y.S., S.L., G.K., Y.G., D.Y.K. Statistical analysis: Y.S., Funding acquisition: F.B.G., R.P.R., P.K. Manuscript preparation – Original draft, review, editing: Y.S., F.B.G., R.P.R., P.K.

## Materials and Methods

### Generation of the plasmid constructs

All oligonucleotides were obtained from Integrated DNA Technologies. Oligonucleotide I-F/R (Supplementary file 1) contains part of a *Hin*dIII site followed by 113 nucleotides that are normally upstream of the G4C2 repeats and then by three G_4_C_2_ repeats. Oligonucleotide II-F/R contains 10 G_4_C_2_ repeats followed by part of a *Not*I site. These two oligonucleotides were phosphorylated, annealed, and then ligated into restriction sites of *Hin*dIII and *Not*I of a pAG plasmid. The plasmid was then digested with HindIII and *Bam*HI. The *Hin*dIII-*Bam*HI fragment was digested with *Ban*II, and the resultant *Hin*dIII-*Ban*II fragment was then ligated with oligonucleotide II-F/R into the pAG plasmid. This approach was repeated three times with similar digestions and ligations of oligonucleotide II. Finally, the *Hin*dIII-*Ban*II fragment was ligated with oligonucleotide III-F/R (which contains 2 G_4_C_2_ repeats followed by a 99 bp flanking sequence and then followed by part of the *Not*I site) into the pAG plasmid (referred to as 113bp-35RG4C2-99bp plasmid). To delete stop codons after the C4G2 repeats, the plasmid was treated with BfaI and NotI, and the digested fragment was ligated with oligonucleotide IV-F/R. To add sequence upstream from the C4G2 repeats, a 543 bp portion (408-950 of NCBI reference sequence, NC_000009.12) of the *C9ORF72* gene from HEK293 genomic DNA was amplified by PCR using the primer shown in Supplementary file 1. The amplified construct was then ligated with the BtgI/NotI-digested fragment of the 113bp-35RG4C2-99bp plasmid into XbaI and NotI sites of pcDNA6/V5-His A plasmid (referred to as 609bp-35RC4G2 plasmid). To further increase the length of sequence upstream from C4G2 repeats, a 392 bp portion (951-1342 of NCBI reference sequence, NC_000009.12) of *C9ORF72* gene from HEK293 genomic DNA was amplified by PCR using the primer shown in Supplementary file 1. The amplified construct was then ligated with the XbaI/NotI fragment of 609bp-35RC4G2 plasmid into HindIII and NotI sites of the pAG plasmid (referred to as AS-C9 plasmid). The ΔC9 plasmid ^21^ was generated as previously described.

To mutate sequences, a 560bp portion upstream from the repeats in the AS-C9 plasmid was amplified by PCR using a primer shown in Supplementary file 1. The amplified portion was then ligated into the HindIII and NotI sites of pcDNA6/V5-His A plasmid. Mutations were made with Q5^®^ Site-Directed Mutagenesis Kit (New England Biolabs) using primer sets (Supplementary file 1). The StuI/BtgI portion of the resultant mutants was then cloned back into the StuI and NotI sites of AS-C9 plasmid with BtgI/NotI portion of AS-C9 plasmid using the primer sets in Supplementary file 1.

### Cell culture

HEK293 and NSC34 cells were cultured in DMEM supplemented with 10% FBS, 2 mM L-Glutamine, 100 U/ml Penicillin and 100 µg/ml Streptomycin.

### Luciferase Assay

The cells were plated in 24-well plates at 5 × 10^4^ per well and then cotransfected using Lipofectamine LTX (Thermo Fisher Scientific) with 100 ng of the plasmid along with 100 ng fLuc plasmid as a transfection control. After 48h, the cells were lysed with 1× passive lysis buffer (Promega). Levels of nLuc and fLuc were assessed with the Nano-Glo Dual-Luciferase Reporter assay system (Promega) and a Wallac 1420 VICTOR 3V luminometer (Perkin Elmer) according to the manufacturer’s protocol.

### Western blotting

The cells were plated in 6-well plates at 2 × 10^5^ per well and then cotransfected with 2.5 µg of plasmids using Lipofectamine LTX (ThermoFisher Scientific). After 48h, cell lysates were prepared using RIPA buffer (50 mM Tris-HCl, pH 7.5; 150 mM NaCl; 0.1% SDS; 0.5% sodium deoxycholate; 5 mM EDTA containing 1 × Halt^TM^ Protease inhibitor Cocktail). Lysates were subjected to electrophoresis on Mini-PROTEAN TGX Gels (BIO-RAD), and then transferred to Amersham Hybond P 0.45 µm PVDF membranes (GE Healthcare). The membrane was blocked with 5% non-fat skim milk in Tris-buffered saline containing 0.05% Tween-20 for 1 h at room temperature, and then incubated overnight at 4 °C with primary antibodies against poly-PR (1:1000, ABN1354, EMD Millipore), poly-GP (1:1000, TALS 828.179, Target ALS), eIF2D (1:1000, 12840-1-AP, Proteintech) and α-tubulin (1:5000, YL1/2, Abcam). Following washing, the membrane was incubated for 1 h at room temperature with anti-mouse (1:5000, GE Healthcare), anti-rabbit (1:5000, GE Healthcare), or anti-rat horseradish peroxidase–conjugated secondary antibodies (1:1000, Cell Signaling Technology). The signal was detected using SuperSignal West Dura Extended Duration Substrate (ThermoFisher Scientific) and analyzed using ChemiDoc MP Imaging System and Image Lab software (version 6.0.1, Bio-Rad).

### Generation of *EIF2D* knockout cells by CRISPR/Cas9 gene editing

A single guide RNA (sgRNA) (GCAGTGACTGTGTACGTGAG) that targets exon 2 of eIF2D was cloned into lentiCRISPR v2 plasmid (Addgene). HEK293 cells were plated into 6-well plates at 4 × 10^5^ cells per well, and then transfected using Lipofectamine LTX with 2.5 µg lentiCRISPR v2 plasmids containing the sgRNA sequence. Transfected cells were selected using 3 µg/ml puromycin for 3 days. *EIF2D* knockout cell clones were obtained by limited dilution. The resulting *EIF2D* knockout cells carry allele-specific mutations, as follows. Compared to the WT GGATGCAGTGACTGTGTACGTGAGTGGTGG sequence, one allele GGATGCAGTGACTGTGTACG**T**TGAGTGGTGG has a single nucleotide insertion shown bolded while the other allele contains a two-nucleotide deletion GGATGCAGTGACTGTGTA— TGAGTGGTGG. Both alleles lead to a premature stop codon, likely resulting in two different truncated eIF2D proteins with the following respective sequence: MFAKAFRVKSNTAIKGSDRRKLRADVTTAFPTLGTDQVSELVPGKEELNIVKLYAHKGDAVT VYEWW and MFAKAFRVKSNTAIKGSDRRKLRADVTTAFPTLGTDQVSELVPGKEELNIVKLYAHKGDAVTVYVEWW.

### Knockdown of eIF2D in HEK293 cells

shRNA plasmids against human eIF2D were prepared using previously published methods ^21^. In brief, oligonucleotides with an siRNA sequence were cloned into the *Bam*HI and *Hin*dIII sites of p*Silencer* 2.1-U6 neo Vector (ThermoFisher Scientific) according to the manufacturer’s protocol. The latter kit also contained a control shRNA vector. For luciferase assays (shown above), the cells were plated in 24-well plates at 5 × 10^4^ per well and cotransfected with 50 ng of the AS-C9 plasmids and 50 ng of the fLuc plasmids along with 500 ng of either control shRNA or anti-eIF2D shRNA using Lipofectamine LTX (ThermoFisher Scientific).

### Motor Neuron Differentiation from human iPSC lines

Human motor neurons were differentiated as previously described from a published iPSC line obtained from a *C9ORF72* carrier (FTD26-6), as well as an isogenic control line that had a CRISPR/Cas9-mediated deletion of expanded GGGGCC repeats^37, 38^. Briefly, iPSCs were plated and expanded in mTSER1 medium (Stem Cell Technologies) in Matrigel-coated wells. Twenty-four hours after plating, the culture medium was replaced every other day with neuroepithelial progenitor (NEP) medium, DMEM/F12 (Gibco), neurobasal medium (Gibco) at 1:1, 0.5X N2 (Gibco), 0.5X B27 (Gibco), 0.1 mM ascorbic acid (Sigma), 1X Glutamax (Invitrogen), 3 µM CHIR99021 (Tocris Bioscience), 2 µM DMH1 (Tocris Bioscience), and 2 µM SB431542 (Stemgent) for 6 days. NEPs were dissociated with accutase, split 1:6 into Matrigel-coated wells, and then cultured for 6 days in motor neuron progenitor induction medium (NEP with 0.1 µM retinoic acid and 0.5 µM purmorphamine, both from Stemgent). Motor neuron progenitors were dissociated with accutase to generate suspension cultures, and the cells were cultured in motor neuron differentiation medium (NEP with 0.5 µM retinoic acid and 0.1 µM purmorphamine). After 6 days, the cultures were dissociated into single cells, and seeded on Matrigel-coated plates in motor neuron medium, 0.5X B27 supplement, 0.1 mM ascorbic acid, 1X Glutamax, 0.1 µM Compound E (Calbiochem), 0.26 µg/ml cAMP, 1 µg/ml Laminin (Sigma), 10 ng/ml GDNF (R&D Systems), and 10 ng/ml GDNF (R&D Systems), and 10 ng/ml BDNF. Motor neurons were cultured for 5 weeks.

### SiRNA Knockdown

After 3 weeks in neuron culture media, motor neurons were transfected with a siRNA specific to *eIF2D* mRNA or a scrambled control. For the transfection, lipofectamine RNAiMAX (ThermoFisher Scientific) was first diluted in Opti-MEM medium, and then both eIF2D and scrambled control siRNAs were separately diluted in Opti-MEM medium at room temperature. Diluted siRNA and diluted lipofectamine RNAiMAX (1:1 ratio) were then mixed and incubated for 20 min. The siRNA-lipid complex solution was then brought up to the appropriate volume with MN culture medium. The culture medium in the plate was aspirated and replaced with a siRNA-lipid complex at a final concentration of 60 pmol siRNA in 1.5 ml medium per 1,000,000 cells. After 24 hours, the medium was replaced with a normal motor neuron medium. This process was repeated two more times at 26 and 31 days in culture. After 36 days in culture, we measured siRNA efficiency and levels of DPRs in harvested motor neurons.

### RNA Extraction and Quantitative Real-time PCR

Total RNA from iPSC-derived motor neurons was extracted with the RNeasy Mini Kit (Qiagen) and then reverse transcribed to cDNA with the TaqMan Reverse Transcription Kit (Applied Biosystems). Quantitative PCR was carried out with SYBR Green Master Mix (Applied Biosystems). Using primers listed in SI Appendix, Table, Ct values for each gene were normalized to actin and GAPDH. Relative mRNA expression was calculated with the double delta Ct method.

### Poly-GR and Poly-GP measurement in iPSC-derived neurons

DPR levels in iPSC-derived neurons were detected using the Meso Scale Discovery (MSD) Immunoassay platform as previously reported^17^. In brief, cells were lysed using Tris based lysis buffer, and lysates were adjusted to equal concentrations and loaded in duplicate wells. Background subtracted electrochemiluminescence (ECL) signals were presented as percentage.

### Soluble and insoluble fractionation for measurement of poly-GA

Motor neurons were lysed in RIPA buffer (Boston BioProducts, BP-115D) with protease and phosphatase inhibitors. The lysates were rotated for 30 min at 4 C, followed by centrifugation at 13,500 rpm for 20 min. The supernatant was removed and used as the soluble fraction. Protein concentrations of the soluble fraction were determined by the BCA assay (Thermo Fisher Scientific, Cat # 23227). To remove carryovers, the pellets were washed with RIPA buffer, and then resuspended in the same buffer with 2% SDS followed by sonication on ice. The lysates were rotated for 30 min at 4C, then spun at 14,800 rpm for 20 min at 4C. The supernatant was removed and used as insoluble fraction. Protein concentrations of the insoluble fraction were determined by Pierce™ 660 nm Protein Assay (Thermo Fisher Scientific, 22660).

### Measurement of poly-GA in iPSC-derived neurons

Poly-GA in soluble and insoluble motor neuron lysates was measured using a Meso Scale Discovery sandwich immunoassay. A human/murine chimeric form of anti-GA antibody chGA3 was used for capture, and a human anti-GA antibody GA4 with a SULFO-tagged anti-human secondary antibody was used for detection. Poly-GA concentrations were interpolated from the standard curve using 60X-GA expressed in HEK 293 cells and presented as percentage. For background correction, values from no-repeats neuron samples were subtracted from the corresponding test samples.

### Statistical analysis

Statistical analysis was performed by one-way ANOVA with Tukey’s multiple comparison test and two-way ANOVA with the Šídák multiple comparison test using GraphPad Prism version 9.3.1. A *P*-value of <0.05 was considered significant. The data are presented as mean ± standard error of the mean.

---

**Supplementary File 1: List of primers used for this study**

## References

1. DeJesus-Hernandez, M. et al. Expanded GGGGCC hexanucleotide repeat in noncoding region of C9ORF72 causes chromosome 9p-linked FTD and ALS. Neuron 72, 245–256 (2011).

2. Renton, A. E. et al. A hexanucleotide repeat expansion in C9ORF72 is the cause of chromosome 9p21-linked ALS-FTD. Neuron 72, 257–268 (2011).

3. McEachin, Z. T., Parameswaran, J., Raj, N., Bassell, G. J. & Jiang, J. RNA-mediated toxicity in C9orf72 ALS and FTD. Neurobiol Dis 145, 105055 (2020).

4. Parameswaran, J., et al. Antisense, but not sense, repeat expanded RNAs activate PKR/eIF2α-dependent integrated stress response in C9orf72 FTD/ALS. bioRxiv, 2022.2006.2006.495030 (2022).

5. Taylor, J. P., Brown Jr, R. H. & Cleveland, D. W. Decoding ALS: from genes to mechanism. Nature 539, 197–206 (2016).

6. Zu, T. et al. RAN proteins and RNA foci from antisense transcripts in C9ORF72 ALS and frontotemporal dementia. Proc Natl Acad Sci U S A 110, E4968–4977 (2013).

7. Gendron, T. F. et al. Antisense transcripts of the expanded C9ORF72 hexanucleotide repeat form nuclear RNA foci and undergo repeat-associated non-ATG translation in c9FTD/ALS. Acta Neuropathol 126, 829–844 (2013).

8. Schmitz, A., Pinheiro Marques, J., Oertig, I., Maharjan, N. & Saxena, S. Emerging Perspectives on Dipeptide Repeat Proteins in C9ORF72 ALS/FTD. Front Cell Neurosci 15, 637548 (2021).

9. Green, K. M. et al. RAN translation at C9orf72-associated repeat expansions is selectively enhanced by the integrated stress response. Nat Commun 8, 2005 (2017).

10. Tabet, R. et al. CUG initiation and frameshifting enable production of dipeptide repeat proteins from ALS/FTD C9ORF72 transcripts. Nat Commun 9, 152 (2018).

11. Boivin, M. et al. Reduced autophagy upon C9ORF72 loss synergizes with dipeptide repeat protein toxicity in G4C2 repeat expansion disorders. EMBO J 39, e100574 (2020).

12. Sonobe, Y. et al. Translation of dipeptide repeat proteins from the C9ORF72 expanded repeat is associated with cellular stress. Neurobiol Dis 116, 155–165 (2018).

13. Lampasona, A., Almeida, S. & Gao, F. B. Translation of the poly(GR) frame in C9ORF72-ALS/FTD is regulated by cis-elements involved in alternative splicing. Neurobiol Aging 105, 327–332 (2021).

14. van ’t Spijker, H. M., et al. Ribosome profiling reveals novel regulation of C9ORF72 GGGGCC repeat-containing RNA translation. RNA 28, 123–138 (2022).

15. Almeida, S. et al. Production of poly(GA) in C9ORF72 patient motor neurons derived from induced pluripotent stem cells. Acta Neuropathol 138, 1099–1101 (2019).

16. Tran, H. et al. Suppression of mutant C9orf72 expression by a potent mixed backbone antisense oligonucleotide. Nat Med 28, 117–124 (2022).

17. Krishnan, G. et al. Poly(GR) and poly(GA) in cerebrospinal fluid as potential biomarkers for C9ORF72-ALS/FTD. Nat Commun 13, 2799 (2022).

18. Cheng, W. et al. C9ORF72 GGGGCC repeat-associated non-AUG translation is upregulated by stress through eIF2α phosphorylation. Nat Commun 9, 51 (2018).

19. Yamada, S. B. et al. RPS25 is required for efficient RAN translation of C9orf72 and other neurodegenerative disease-associated nucleotide repeats. Nat Neurosci 22, 1383–1388 (2019).

20. Ayhan, F. et al. SCA8 RAN polySer protein preferentially accumulates in white matter regions and is regulated by eIF3F. EMBO J 37, (2018).

21. Sonobe, Y. et al. A C. elegans model of C9orf72-associated ALS/FTD uncovers a conserved role for eIF2D in RAN translation. Nat Commun 12, 6025 (2021).

22. Green, K. M., Miller, S. L., Malik, I. & Todd, P. K. Non-canonical initiation factors modulate repeat-associated non-AUG translation. Hum Mol Genet, In print (2022).

23. Lee, K. H. et al. C9orf72 Dipeptide Repeats Impair the Assembly, Dynamics, and Function of Membrane-Less Organelles. Cell 167, 774–788.e717 (2016).

24. Lin, Y. et al. Toxic PR Poly-Dipeptides Encoded by the C9orf72 Repeat Expansion Target LC Domain Polymers. Cell 167, 789–802.e712 (2016).

25. Kwon, I. et al. Poly-dipeptides encoded by the C9orf72 repeats bind nucleoli, impede RNA biogenesis, and kill cells. Science 345, 1139–1145 (2014).

26. Rudich, P. et al. Nuclear localized C9orf72-associated arginine-containing dipeptides exhibit age-dependent toxicity in C. elegans. Hum Mol Genet 26, 4916–4928 (2017).

27. Wen, X. et al. Antisense proline-arginine RAN dipeptides linked to C9ORF72-ALS/FTD form toxic nuclear aggregates that initiate in vitro and in vivo neuronal death. Neuron 84, 1213–1225 (2014).

28. Maor-Nof, M. et al. p53 is a central regulator driving neurodegeneration caused by C9orf72 poly(PR). Cell 184, 689–708.e620 (2021).

29. Zhang, Y. J. et al. Heterochromatin anomalies and double-stranded RNA accumulation underlie C9orf72 poly(PR) toxicity. Science 363, eaav2606 (2019).

30. Hao, Z. et al. Motor dysfunction and neurodegeneration in a C9orf72 mouse line expressing poly-PR. Nat Commun 10, 2906 (2019).

31. Gleason, A. C., Ghadge, G., Chen, J., Sonobe, Y. & Roos, R. P. Machine learning predicts translation initiation sites in neurologic diseases with nucleotide repeat expansions. PLoS One 17, e0256411 (2022).

32. Chew, J. et al. Aberrant deposition of stress granule-resident proteins linked to C9orf72-associated TDP-43 proteinopathy. Mol Neurodegener 14, 9 (2019).

33. Mizielinska, S. et al. C9orf72 repeat expansions cause neurodegeneration in Drosophila through arginine-rich proteins. Science 345, 1192–1194 (2014).

34. Freibaum, B. D. et al. GGGGCC repeat expansion in C9orf72 compromises nucleocytoplasmic transport. Nature 525, 129–133 (2015).

35. Gendron, T. F. et al. Poly(GP) proteins are a useful pharmacodynamic marker for C9ORF72-associated amyotrophic lateral sclerosis. Sci Transl Med 9, eaai7866 (2017).

36. Lehmer, C. et al. Poly-GP in cerebrospinal fluid links C9orf72-associated dipeptide repeat expression to the asymptomatic phase of ALS/FTD. EMBO Mol Med 9, 859–868 (2017).

37. Lopez-Gonzalez, R. et al. Partial inhibition of the overactivated Ku80-dependent DNA repair pathway rescues neurodegeneration in C9ORF72-ALS/FTD. Proc Natl Acad Sci U S A 116, 9628–9633 (2019).

38. Lopez-Gonzalez, R. et al. Poly(GR) in C9ORF72-Related ALS/FTD Compromises Mitochondrial Function and Increases Oxidative Stress and DNA Damage in iPSC-Derived Motor Neurons. Neuron 92, 383–391 (2016).

39. Cheng, W. et al. CRISPR-Cas9 Screens Identify the RNA Helicase DDX3X as a Repressor of C9ORF72 (GGGGCC)n Repeat-Associated Non-AUG Translation. Neuron 104, 885–898.e888 (2019).

40. Goodman, L. D. et al. eIF4B and eIF4H mediate GR production from expanded G4C2 in a Drosophila model for C9orf72-associated ALS. Acta Neuropathol Commun 7, 62 (2019).

41. Richter, J. D. & Sonenberg, N. Regulation of cap-dependent translation by eIF4E inhibitory proteins. Nature 433, 477–480 (2005).

42. Kramer, N. J. et al. Spt4 selectively regulates the expression of C9orf72 sense and antisense mutant transcripts. Science 353, 708–712 (2016).

43. Chintalaphani, S. R., Pineda, S. S., Deveson, I. W. & Kumar, K. R. An update on the neurological short tandem repeat expansion disorders and the emergence of long-read sequencing diagnostics. Acta Neuropathol Commun 9, 98 (2021).

44. Depienne, C. & Mandel, J. L. 30 years of repeat expansion disorders: What have we learned and what are the remaining challenges? Am J Hum Genet 108, 764–785 (2021).

45. Gao, F. B., Richter, J. D. & Cleveland, D. W. Rethinking Unconventional Translation in Neurodegeneration. Cell 171, 994–1000 (2017).

